# Single-cell RNA-seq reveals lineage-specific regulatory changes of fibroblasts and vascular endothelial cells in keloid

**DOI:** 10.1101/2020.05.14.095323

**Authors:** Xuanyu Liu, Wen Chen, Meng Yuan, Zhujun Li, Tian Meng, Jie Chen, Nanze Yu, Xiao Long, Zhou Zhou

**Author notes:** X. Liu and W. C. contribute equally to this manuscript. Correspondence author; email: Z.Z. and X. Long.

## Abstract

Keloid is a benign dermal fibrotic disorder with some features similar to malignant tumors such as hyper-proliferation, apoptosis resistance and invasion. keloid remains a therapeutic challenge in terms of high recurrence rate and lack of satisfactory medical therapies, which is partially due to the incomplete understanding of keloid pathogenesis. A thorough understanding of the cellular and molecular mechanism of keloid pathogenesis would facilitate the development of novel medical therapies for this disease. Here, we performed single-cell RNA-seq of 28,064 cells from keloid skin tissue and adjacent relatively normal tissue. Unbiased clustering revealed substantial cellular heterogeneity of the keloid tissue, which included 21 cell clusters assigned to 11 cell lineages. Differential proportion analysis revealed significant expansion for fibroblasts and vascular endothelial cells in keloid compared with control, reflecting their strong association with keloid pathogenesis. We then identified five previously unrecognized subpopulations of keloid fibroblasts and four subpopulations of vascular endothelial cells. Comparative analyses were performed to identify the dysregulated pathways, regulators and ligand-receptor interactions for keloid fibroblasts and vascular endothelial cells, the two important cell lineages in keloid pathogenesis and for medical interventions. Our results highlight the roles of transforming growth factor beta and Eph-ephrin signaling pathways in both the aberrant fibrogenesis and angiogenesis of keloid. Critical regulators and signaling receptors implicated in the fibrogenesis of other fibrotic disorders, such as *TWIST1, FOXO3, SMAD3* and *EPHB2*, ranked at the top in the regulatory network of keloid fibroblasts. In addition, tumor-related pathways such as negative regulation of *PTEN* transcription were found to be activated in keloid fibroblasts and vascular endothelial cells, which may be responsible for the malignant features of keloid. Our study put novel insights into the pathogenesis of keloid, and provided potential targets for medical therapies. Our dataset also constitutes a valuable resource for further investigations of the mechanism of keloid pathogenesis.

## Introduction

Keloid is a dermal fibrotic disorder following a aberrant wound healing response (Glass, 2017). Histologically, keloid scars are characterized by excessive extracellular matrix (ECM) deposition and rich vasculature (Ashcroft et al., 2013). Despite that keloid is classified as a benign dermal growth, it demonstrates biological features similar to malignant tumors such as hyper-proliferation, apoptosis resistance and invasion (Mari et al., 2015). Keloid scars grow beyond the boundaries of the original wound, causing pain, pruritus and contracture which leads to serious physical and psychological burden for patients (Gauglitz et al., 2011). Keloid is common and has a higher prevalence in Asians and Africans, especially for dark-skinned individuals (Zhu et al., 2013). Although a wide range of therapies currently being used, keloid remains a therapeutic challenge in terms of high recurrence rate and lack of satisfactory medical therapies (Mari et al., 2015). This is partially due to the incomplete understanding of keloid pathogenesis, although both environmental and genetic factors have been implicated (Nakashima et al., 2010; Shih et al., 2010). A thorough understanding of the cellular and molecular mechanism of keloid pathogenesis would facilitate the development of novel medical therapies for this disease.

Recent technical advances in single-cell RNA-seq have enabled the transcriptomes of tens of thousands of cells to be assayed at a single-cell resolution (Zheng et al., 2017). Compared with the average expression of genes from a mixed cell population obtained in bulk RNA-seq, large-scale single-cell RNA-seq allows unbiased cellular heterogeneity dissection and regulatory network construction at an unprecedented scale and resolution (Kulkarni et al., 2019). Single-cell RNA-seq is therefore emerging as a powerful tool for understanding the cellular and molecular mechanism of the pathogenesis in a variety of diseases including fibrotic disorders such as pulmonary fibrosis (Reyfman et al., 2019) and lupus nephritis (Der et al., 2019). Single-cell RNA-seq have already been applied to dissect the cellular heterogeneity of human skin in normal states (Philippeos et al., 2018) and diseased conditions, such as atopic dermatitis (He et al., 2020) and inflamed epidermis (Cheng et al., 2018). Previous efforts have been made to examine the transcriptomic alterations in keloid tissue using bulk RNA-seq or microarray (Liang et al., 2015; Onoufriadis et al., 2018; Wang et al., 2019). However, to our knowledge, the cellular heterogeneity and regulatory changes in keloid have not yet been systematically investigated at single-cell resolution.

In this study, we performed single-cell RNA-seq of 28,064 cells from keloid skin tissue and adjacent relatively normal tissue. Comparative analyses were performed to identify the dysregulated pathways, regulators and ligand-receptor interactions for keloid fibroblasts and vascular endothelial cells, the two important cell lineages in keloid pathogenesis and for medical interventions. Our study put novel insights into the pathogenesis of keloid, and provided potential targets for medical therapies. Our dataset also constitutes a valuable resource for further investigations of the mechanism of keloid pathogenesis.

## Results

### Single-cell RNA-seq reveals cellular diversity and heterogeneity of keloid skin tissue

To dissect the cellular heterogeneity and explore the regulatory changes of keloid skin tissue, we sampled keloid lesional skin tissue (CASE) and matched relatively normal tissue adjacent to the keloid scar (CTRL) from four patients (female; Chinese; 26-32 years old; lesion position: chest; Table S1). The eight samples were dissociated to single cells and subjected to single-cell RNA-seq (Figure 1A). After stringent quality control, we obtained transcriptomes of 28,064 cells (CASE: 12,425; CTRL:15,639). Unbiased clustering revealed 21 cell clusters (Figure 1B). Based on hierarchical clustering (Figure 1C) and established lineage-specific marker genes (Figure 1D), we assigned these clusters into 11 cell lineages. The keratinocyte lineage (marked by *KRT5* and *KRT14*) (Joost et al., 2016), including the cluster c0, c1, c8, c10, c11 and c15, accounted for the largest proportion (41.7%) of cells. Vascular endothelial cells (marked by *PECAM1* and *CDH5*) (Kalucka et al., 2020), including the cluster c2, c4, c5 and c18, accounted for the second largest proportion (26.3%) of cells. Lymphatic endothelial cells (c16) differed from vascular endothelial cells by expressing lineage-specific markers such as *LYVE1* and *PROX1* (Johnson et al., 2008). The fibroblast lineage (marked by *PDGFRA* and *DCN*) (Guerrero-Juarez et al., 2019) had two clusters c3 and c9. In addition, we found other typical skin lineages including sweat gland cells (marked by *AQP5* and *MUCL1*) (He et al., 2020), smooth muscle cells (marked by *MYH11* and *CNN1*) (Liu et al., 2019), mural cells (marked by *PDGFRB* and *RGS5*) (Holm et al., 2018), neural cells (marked by *NRXN1* and SCN7A) (He et al., 2020), melanocyte (marked by *MLANA* and *DCT*) (Miller et al., 2004) and Schwann cells (marked by *GLDN* and *TJP1*) (He et al., 2020). These clusters showed distinct molecular signatures (Figure 1E; Table S2), reflecting the cellular diversity and heterogeneity of keloid skin tissues.

**Figure 1.**
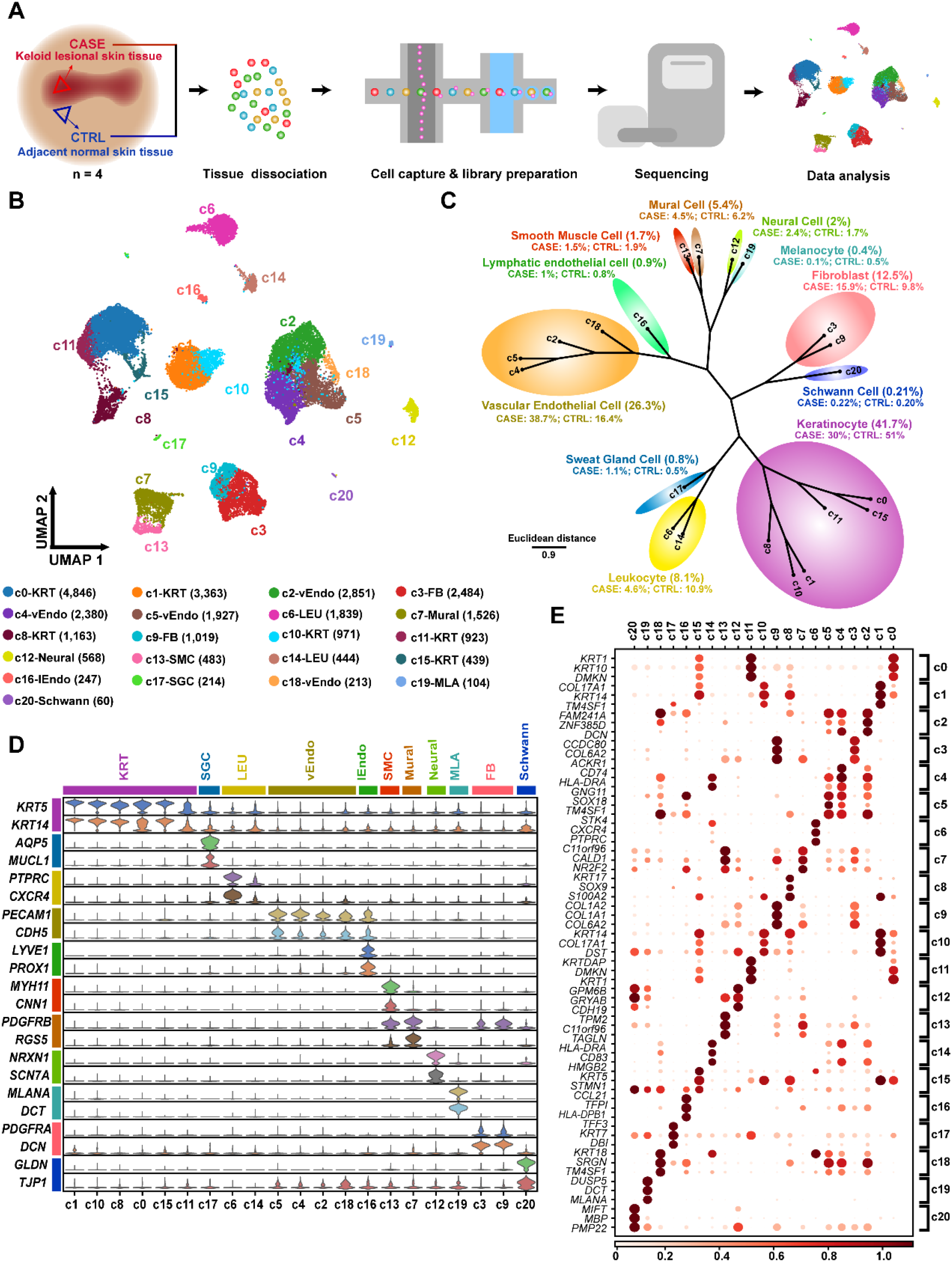
Single-cell RNA-seq reveals cellular diversity and heterogeneity of keloid skin tissue. **(A)**Schematic representation of the experimental procedure. Keloid lesional (CASE) and adjacent normal (CTRL) skin tissues were harvested separately at the time of surgery (n=4). **(B)** Unbiased clustering of 28,064 cells reveals 21 cellular clusters. Clusters are distinguished by different colors. The number in the parenthesis means the count of cells. **(C)** Hierarchical clustering of the clusters based on the average expression of 2,000 most variable genes. **(D)** Expression of the established marker genes for each lineage in each cluster. **(E)** Representative molecular signatures for each cell cluster. The area of the circles indicates the proportion of cells expressing the gene, and the color intensity reflects the expression intensity. FB: fibroblast; KTR: Keratinocyte; LEU: Leukocyte; lEndo: lymphatic endothelial cell; MLA: Melanocyte; SGC: sweat gland cell; SMC: smooth muscle cell; vEndo: vascular endothelial cell

### Differential proportion analysis reveals significant expansion for fibroblasts and vascular endothelial cells in keloid compared with relatively normal skin tissue

We next tried to identify keloid-associated cell lineages or clusters, which significantly expanded or contracted in CASE versus CTRL. Visualization for cellular density revealed dramatic changes in relative proportion for multiple cell lineages (Figure 2A). For example, lineage expansion was observed for vascular endothelial cells and fibroblasts in keloid tissue, while contraction was observed for keratinocytes and leukocytes. We further performed statistical tests, which took individual patient into account (Figure S1). Only the vascular endothelial cells reached statistical significance (one-way paired t-test p-value=0.01; Figure 2B). Similarly, we performed statistical tests at the cluster level (Figure 2C). Three vascular endothelial clusters, c4, c5 and c18, significantly expanded in CASE. Notably, one fibroblast cluster, c9, also significantly expanded. Taken together, differential proportion analysis revealed significant expansion for fibroblasts and vascular endothelial cells in keloid compared with relatively normal skin tissue, reflecting their strong association with keloid pathogenesis. Our study thus focused on these two cell lineages.

**Figure 2.**
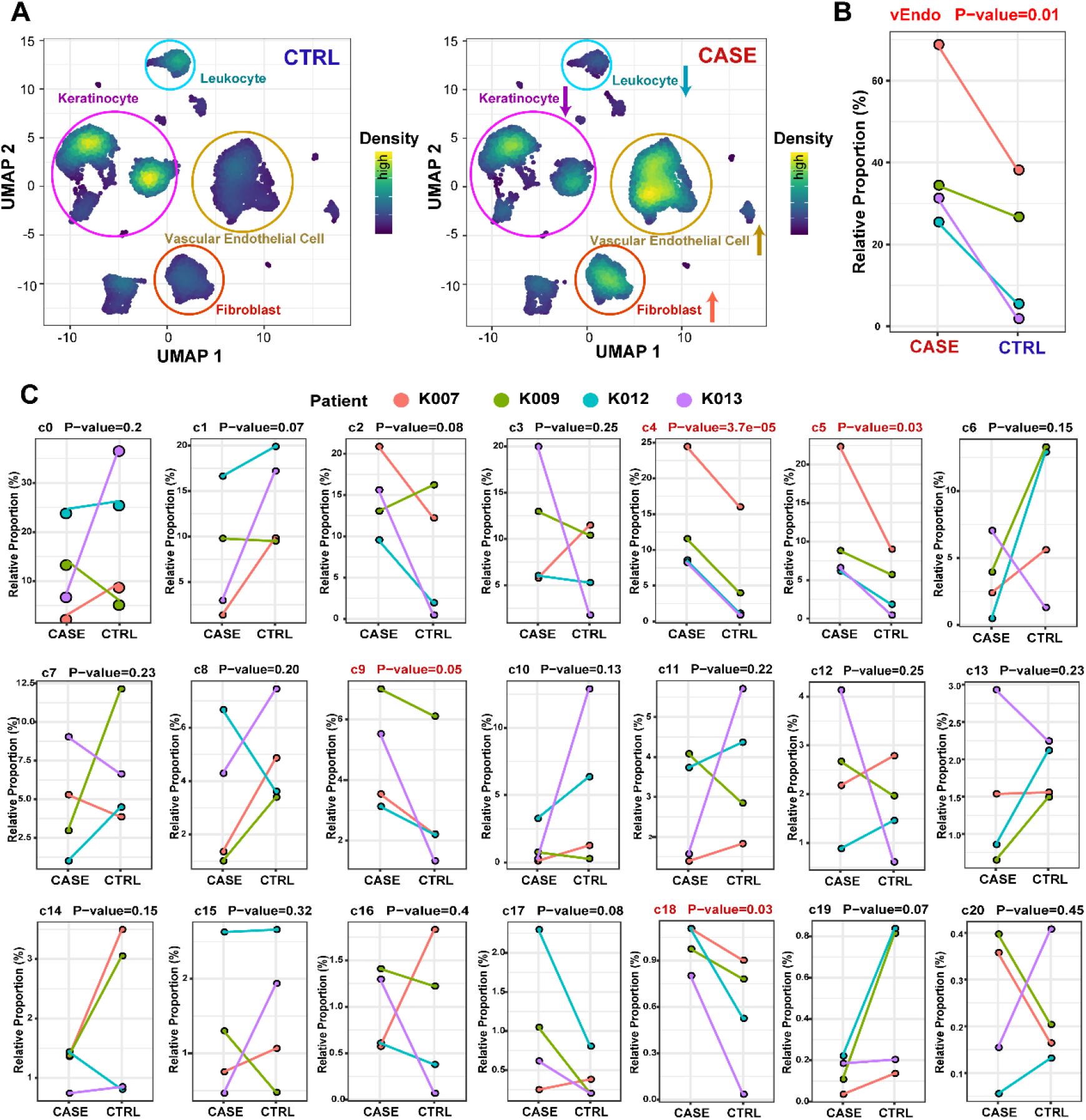
Differential proportion analysis reveals significant lineage expansion for fibroblasts and vascular endothelial cells in keloid compared with relatively normal skin tissue. **(A)** Visualization for cellular density reveals dramatic changes in proportion for multiple cell lineages in CASE versus CTRL. Cells are randomly sampled for equal number of cells in CASE (n=12,425) and CTRL (n=12,425) in this analysis. **(B)** Significant expansion of the vascular endothelial lineage in CASE versus CTRL for each of the patients. **(C)** Only the vascular endothelial clusters c4, c5 and c18 as well as the fibroblast cluster c9 are significantly expanded in CASE versus CTRL for each of the patients. A significant threshold of p-value < 0.05 for one-way paired t-tests was used in B and C.

### Gene set enrichment analysis revealed fibroblast-specific dysregulated pathways in keloid versus normal skin tissue

Single-cell RNA-seq allows to analyze lineage-specific transcriptomic changes without cell sorting. We identified fibroblast-specific differentially regulated pathways in keloid versus normal skin tissue (Figure 3A; Table S3) through gene set enrichment analysis (GSEA), which facilitates biological interpretation by robustly detecting concordant differences at the gene set or pathway level (Emmert-Streib and Glazko, 2011). Consistent with the excessive ECM deposition in keloid, ECM-related pathways such as extracellular matrix organization, collagen formation and elastic fiber formation were significantly up-regulated (Figure 3A; GSEA FDR q-value < 0.05). Carbohydrate metabolism pathways such as glycosaminoglycan (GAG) metabolism, chondroitin sulfate (CS) biosynthesis and keratan sulfate (KS) biosynthesis were significantly up-regulated (Figure 3A), reflecting metabolic reprogramming of fibroblasts to achieve and sustain the functional state in keloid. This result is consistent with previous reports that CS and Hyaluronic acid (HA), two forms of GAG, were over-accumulated in keloid tissue and related medical treatments would improve keloid pathology (Alaish et al., 1995; Ishiko et al., 2013; Katayama et al., 2020)(Ishiko et al., 2013; Katayama et al., 2020). In agree with the hypoxia, mechanical and oxidative stress in the microenvironment of keloid tissue (Nangole and Agak, 2019), pathways related to cellular responses to stress were also up-regulated (Figure 3A). Notably, several signal transduction pathways were activated in keloid fibroblasts (Figure 3B). The role of transforming growth factor (TGF) beta and canonical WNT signaling pathways have been well established in keloid or other fibrotic disorders with excessive fibrosis (Piersma et al., 2015). In line with this, we observed activation of these two master pathways (Figure 3B). Platelet derived growth factor (PDGF) signaling pathway was significantly activated and genes encoding PDGF receptors such as *PDGFRA* and *PDGFRB* were up-regulated in keloid fibroblasts, which is concordant with the previous observation that fibroblasts from keloid tissue were more responsive to PDGF as compared with normal skin fibroblasts (Haisa et al., 1994). We also found aberrant NOTCH1 signaling activation, which has been implicated in the development of organ fibrosis (Hong et al., 2019). *PTEN*, a known tumor suppressor, is a major growth signaling inhibitor that controls cell growth, survival and proliferation (Chalhoub and Baker, 2009). Intriguingly, we found that a set of genes involved in negative regulation of *PTEN* gene transcription were up-regulated in keloid fibroblasts (Figure 3B), which agrees with the malignant features of keloid fibroblasts such as excessive proliferation and resistance to apoptosis (Lim et al., 2006). Notably, Eph-ephrin signaling pathway, which has recently been recognized to be associated with cardiac fibrosis (Su et al., 2017), were significantly activated in keloid fibroblasts (Figure 3B). Multiple genes encoding the ligands and receptors of Eph-ephrin signaling such as *EFNB1, EFNB2, EPHB2, EPHB3* and *EPHA3* were up-regulated in keloid fibroblasts.

**Figure 3.**
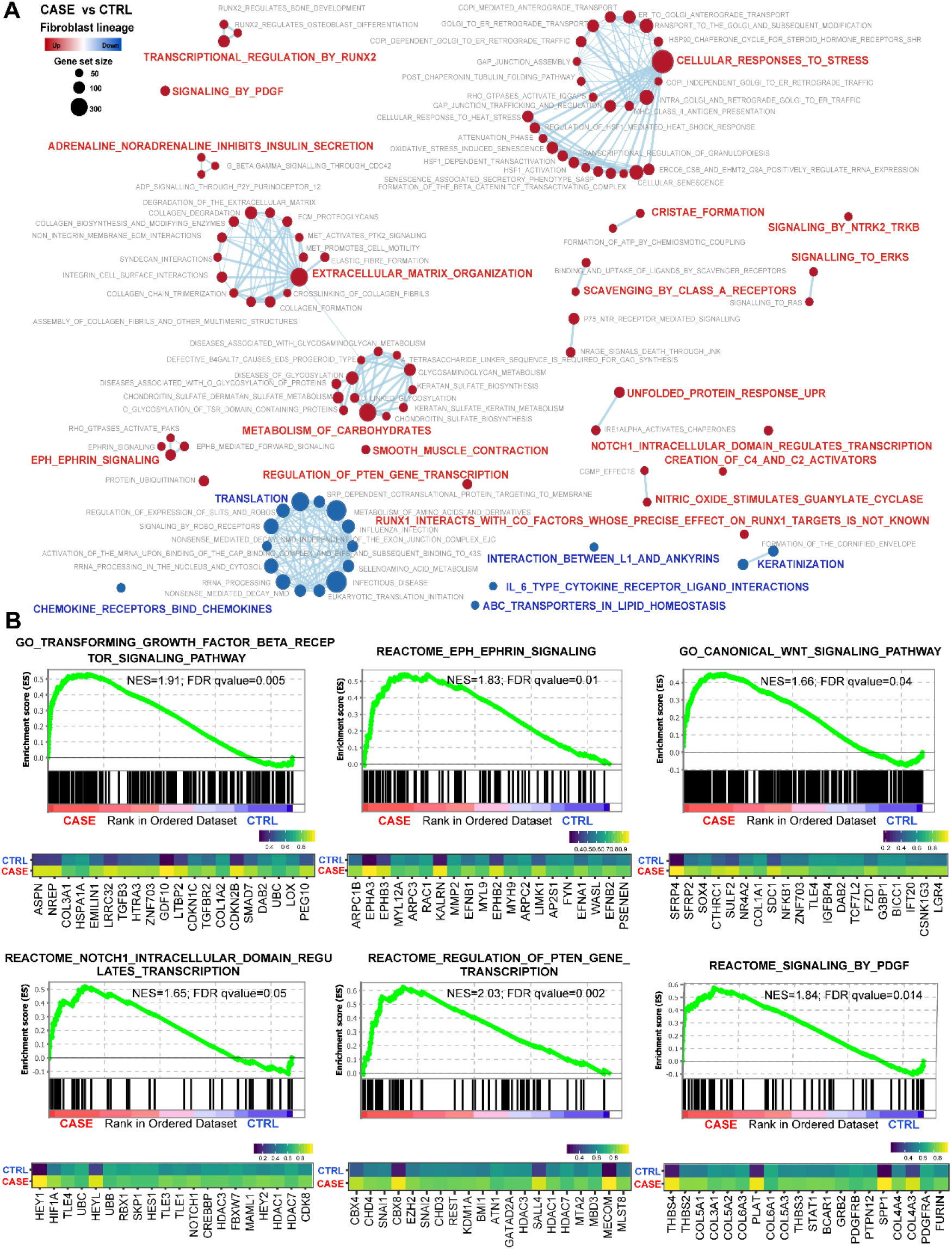
Gene set enrichment analysis reveals fibroblast-specific dysregulated pathways in keloid versus normal skin tissue. **(A)** Network view of differentially regulated REACTOME pathways in keloid fibroblasts. The size of the circle reflects the size of the gene set. The circles in red denote up-regulated pathways, and circles in blue represent down-regulated pathways. A significant threshold of FDR q-value of 0.05 was used. **(B)** Enrichment plots (upper panel) and leading-edge gene expression heatmaps (the top 20 genes; lower panel) for representative signaling pathways up-regulated in keloid fibroblasts. NES: Normalized enrichment score, used to compare analysis results across gene sets. The vertical lines in the enrichment plot show where the members of the gene set appear in the ranked list of genes. Leading-edge genes: a subset of genes in the gene set that contribute most to the enrichment. Average expressions across cells in each group are shown in the heatmap.

### Previously unrecognized cellular heterogeneity of keloid fibroblasts revealed by single-cell transcriptomic data

We next dissected the cellular heterogeneity and characterized the subpopulations of fibroblasts. The fibroblast clusters c3 and c9 showed distinct expression profiles (Figure 4A). Compared with c3, c9 displays more obvious phenotypes of myofibroblasts. For example, significantly higher expression of contractile genes (e.g., *ACTA2, TAGLN* and *MYL9*), collagen and elastin genes (e.g., *COL1A2, COL3A1* and *ELN*) as well as myofibroblast markers (e.g., *FN1* and *CTHRC1*) (Figure 4B; adjusted P-value < 0.01). This result agrees with the significant expansion of c9 in keloid (Figure 2C), reflecting the elevated myofibrogenesis of this fibrotic disorder. To further dissected the cellular heterogeneity, we performed secondary clustering and found five subpopulations (Figure 4C), which had distinct molecular signatures (Figure 4D; Table S4), reflecting their specific functional properties. These fibroblast subpopulations could be marked with specific markers: s0-*PDGFRA*^+^ *POSTN* ^high^, s1-*PDGFRA*^+^ *ABCA8* ^high^, s2-*PDGFRA*^+^ *REL* ^high^, s3-*PDGFRA*^+^ *RGS2* ^high^ and s4-*PDGFRA*^+^ *WISP2* ^high^ (Figure 4E). Correlation analysis was performed to infer the relationships between them (Figure 4F). For example, the closest subpopulation to s0 is s4. Functional enrichment analysis revealed that the signature genes of s0 and s4 were both enriched with ECM-related terms, thus representing two myofibroblast subpopulations (Figure 4G). However, they differed in the components of ECM: s0 with higher level of collagens and s4 with higher level of elastin (Figure 4D). The signature genes of s1 were enriched with terms related with immune response, such as regulation of complement activation and lymphocyte chemotaxis. The signature genes of s2 were enriched with terms related with leukocyte chemotaxis, carbohydrate metabolism and stress response. The signature genes of s3 were enriched with terms related with stress response and differentiation.

**Figure 4.**
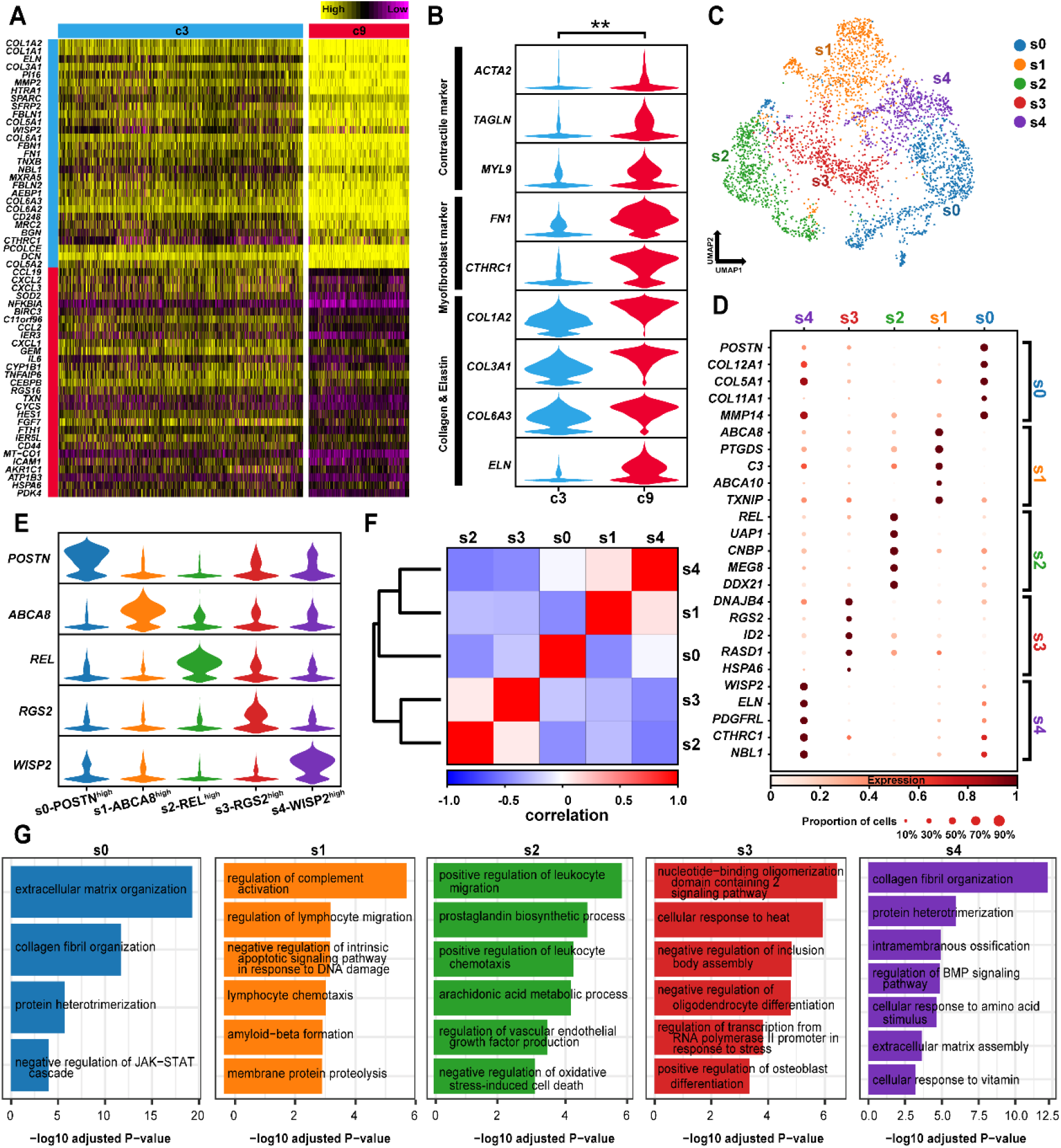
Cellular heterogeneity of keloid fibroblasts revealed by single-cell transcriptomic data. **(A)** Heatmap showing distinct expression profiles between the fibroblast clusters c3 and c9. **(B)** Compared with c3, c9 displays more obvious phenotypes of myofibroblasts. **: adjusted P-value < 0.01. **(C)** Secondary clustering of fibroblasts further identifies five subpopulations. **(D)** The five fibroblast subpopulations display distinct expression profiles. **(E)** Molecular signatures for the five fibroblast subpopulations. **(F)** Correlations among the five subpopulations. **(G)** Functional enrichment for each of the five subpopulations. Adjusted P-value < 0.05.

### Pseudo-temporal ordering of fibroblasts reveals a branched trajectory with a significant shift towards myofibroblast phenotype in keloid

The major source of myofibroblasts are local fibroblasts in wound healing and remodeling (Darby et al., 2014). To further explore the relationships of the fibroblast subpopulations and study the regulatory dynamics during fibroblast-to-myofibroblast phenotypic transition, we performed pseudo-temporal ordering of all fibroblasts with Monocle2 (Qiu et al., 2017). This analysis revealed a branched trajectory with two major branches, i.e., cell fate 1 and cell fate 2 (Figure 5A). The subpopulation s0 and s4 constituted the large majority of cell fate1, which thus represents cellular states of myofibroblast phenotype (Figure 5B). The subpopulation s1 and s3 constituted the large majority of cell fate 2, which thus represents special states of cells associated with immune response, stress response and differentiation. The subpopulation s2 accounted for the largest proportion of the “pre-branch”, which represents the initial states of fibroblasts. Notably, compared with the normal control, the trajectory in keloid displayed a significant shift towards the myofibroblast phenotype (cell fate 1 cells accounted for 28% in CTRL versus 40% in CASE; Figure 5C). Through branched expression analysis modeling (BEAM) tests, we obtained the expression dynamics of 1,204 branch-dependent genes during the cellular state transition from the “pre-branch” to cell fate 1 and cell fate 2 (Figure 5D; Table S5; q-value < 1E-04). Hierarchical clustering of these genes revealed five gene modules, and the reprehensive transcription factors for each gene module were shown in Figure 5D. Among them, the majority of the genes in module I had high expression levels in the ‘pre-branch’ cells. Notably, *TWIST1* is a representative transcription factor of module I, which has recently been recognized as an important pro-fibrotic regulator in a variety of organ fibrotic disorders including skin fibrosis (Ning et al., 2018). In addition, the gene module III and IV had high expression levels in fate 2 and fate 1 cells.

**Figure 5.**
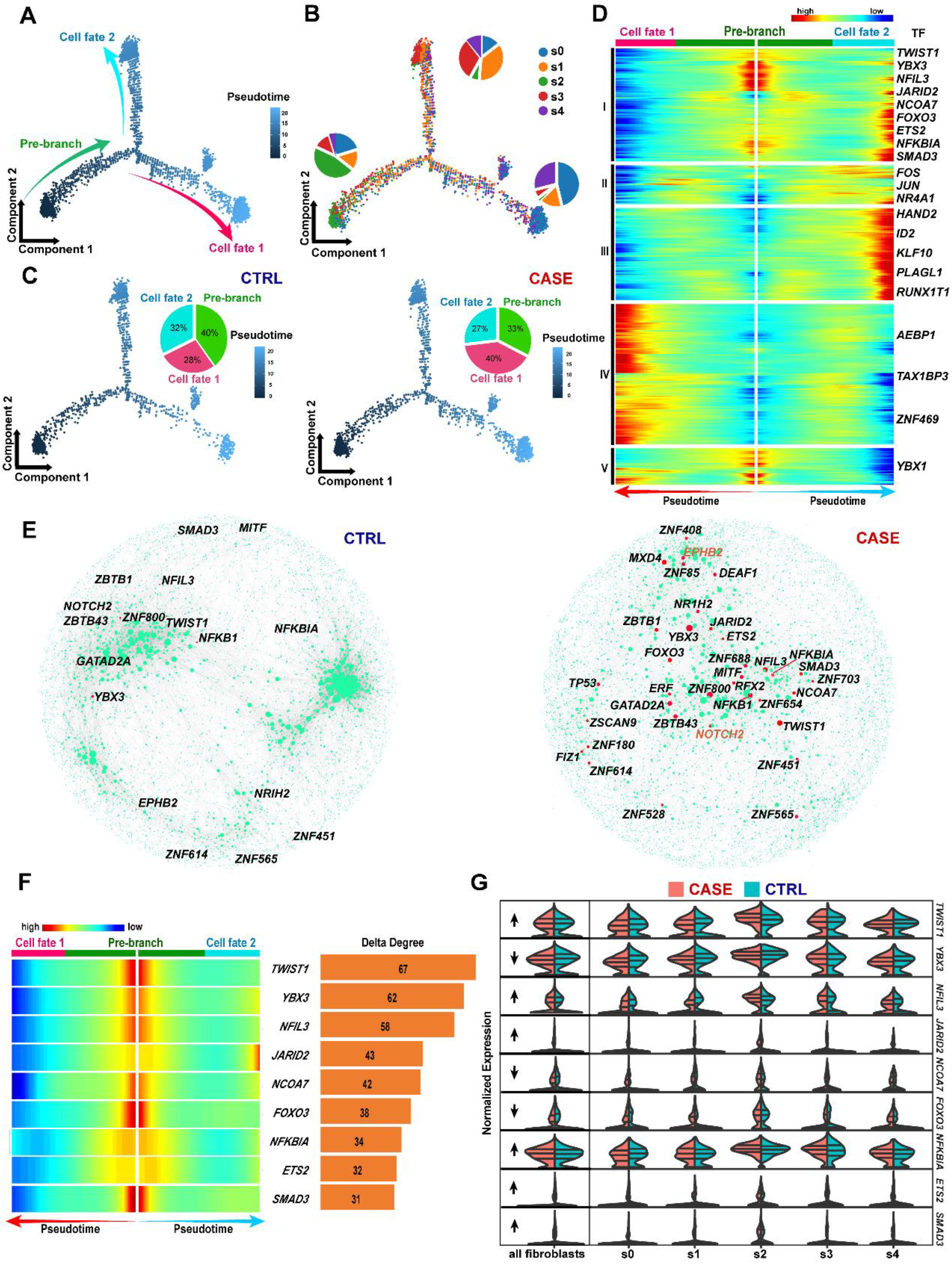
Pseudo-temporal ordering and gene regulatory network analysis of keloid fibroblasts. **(A)** Pseudo-temporal ordering of keloid fibroblasts reveals a branched trajectory. **(B)** Distribution of the five subpopulations in each of the three branches. **(C)** Significant shift towards myofibroblast phenotype (cell fate 1) in CASE (right panel) versus CTRL (left panel). **(D)** Hierarchical clustering of the branch-dependent genes reveals five gene modules. A significant threshold was set to be a q-value of branched expression analysis modeling (BEAM) test < 1E-04. Representative transcription factors are shown. **(E)** Comparative analysis of the gene regulatory networks of fibroblasts between CASE (right panel) and CTRL (left panel) reveals dysregulated genes in keloid fibroblasts. The node size reflects the degree centrality. The transcription factors (in black) and receptors (in orange) in the top 100 genes dysregulated in CASE ranked by delta degree are shown. **(F)** The nine transcription factors that are both branch-dependent and dysregulated in keloid network are all from gene module I, most of which are highly expressed in the “pre-branch”**. (G)** Split violin plot showing the expression of the nine transcription factors in CASE and CTRL.

### Comparative analysis of the gene regulatory networks of fibroblasts between keloid and normal skin tissue reveals dysregulated genes in keloid fibroblasts

To further prioritize transcription factors obtained above, we built gene regulatory networks from single-cell data using a novel method implemented in bigScale2 (Iacono et al., 2019), which allows to quantify biological importance of genes and find key regulators changed in diseased conditions. Figure 5E shows the regulatory networks constructed for fibroblasts from normal and keloid skin tissue. Comparative analysis between the networks of CASE and CTRL revealed a group of genes greatly altered in degree centrality (the number of edges afferent to a given node; Table S6). We focused on the transcription factors and signal receptors in the top 100 genes dysregulated in keloid, which were indicated in Figure 5E and 5F. Notably, among the transcription factor genes, *TWIST1* ranked at the top according to the degree centrality (degree in CASE: 77, degree in CASE: 10), supporting a key role of this regulator in keloid pathogenesis. *EPHB2*, ranked at the top among the signal receptor genes (degree in CASE: 49, degree in CASE: 9), consolidating our view that Eph-ephrin signaling pathway may play critical roles in keloid pathogenesis. We next tried to find transcription factors that were both branch-dependent (Figure 5D) and dysregulated in the keloid network (Figure 5E), which would be more important and reliable given independent analysis approaches. We ultimately found nine transcription factors including *TWIST1, YBX3, NFIL3, JARID2, NCOA7, FOXO3, NFKBIA, ETS2* and *SMAD3* (Figure 5F). Among them, *SMAD3* is a known regulator of keloid through which the TGF beta signaling exerts its pro-fibrotic effects (Darby et al., 2014). *TWIST1* and *FOXO3* have been implicated in fibrogenesis and organ fibrosis (Al-Tamari et al., 2018; Ning et al., 2018). Intriguingly, we found that all the nine candidate regulators were from the gene module I, most of which were highly expressed in the “pre-branch” (Figure 5F). We next examined the expression of these regulators in fibroblasts from keloid and normal tissues (Figure 5G). We found the expression modes of these genes agreed well with that reported in other fibrotic disorders. For example, *TWIST1* and *SMAD3* were up-regulated, and *FOXO3* was down-regulated in keloid versus normal fibroblasts. Taken together, we found previously unrecognized key regulators and signaling receptors that may play critical roles in keloid pathogenesis.

### Cell-cell communication analysis reveals ligand-receptor interaction changes specific to fibroblasts in keloid

The single-cell dataset provided us a unique chance to analyze cell-cell communication mediated by receptor-ligand interactions. To define the cell-cell communication landscape and find its alterations in keloid, we performed analysis using CellPhoneDB 2.0 (Efremova et al., 2019), which contains a curated repository of ligand-receptor interactions and a statistical framework for predicting enriched interactions between two cell types from single-cell transcriptomics data. We found a dense communication network among fibroblasts, vascular endothelial cells, neural cells and keratinocytes in both normal (Figure 6A left panel) and diseased conditions (Figure 6A right panel), which constituted a core signaling module in skin. Notably, the communication network in the normal condition was dominated by vascular endothelial cells, whereas the network in keloid was dominated by fibroblasts, reflecting the role of fibroblasts in keloid pathogenesis. In both conditions, the most abundant interactions occurred between fibroblasts and vascular endothelial cells (Figure 6B), the two most important lineages in keloid pathogenesis. Furthermore, we identified the ligand-receptor pairs shown significant changes in specificity between any one of the non-fibroblast lineages and fibroblasts in keloid versus normal conditions (fibroblasts express receptors and received ligand signals from other lineages; Table S7). As shown in Figure 6C, the significantly altered signals included a wealth of fibrosis-related signals such as TGFB1 signaling, NOTCH1 signaling and fibroblast growth factor (FGF) signaling. Notably, TGFB1, a key ligand of TGF beta signaling pathway implicated in keloid formation (Peltonen et al., 1991), was found to be secreted by lymphatic endothelial cells, leukocytes, mural cells, neural cells, smooth muscle cells and vascular endothelial cells in keloid. The TGFB1-TGF beta receptor 2 interactions between these cell lineages and fibroblasts become significantly more specific in keloid compared with normal conditions (permutation test p-value < 0.05). In addition, we also explored the alterations of ligand signals broadcast by fibroblasts (Figure 6D). Notably, fibroblasts may affect other cells in keloid through alterations in ligand-receptor interactions of the NOTCH signaling, including NVO-NOTCH1, JAG1-NOTCH1, JAG1-NOTCH2 and JAG1-NOTCH4 interactions.

**Figure 6.**
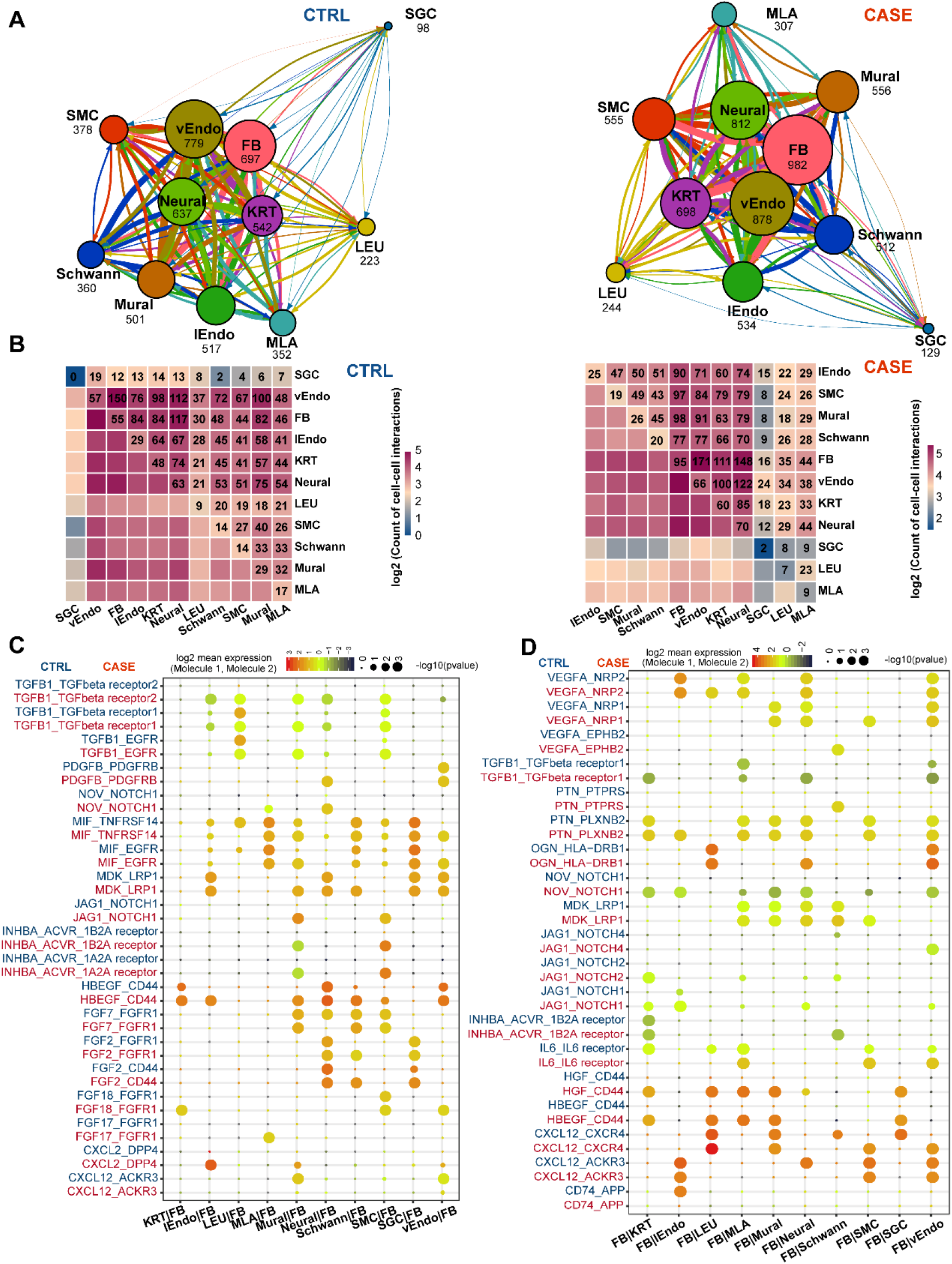
Cell-cell communications among the cell lineages in keloid and relatively normal skin tissues. **(A)** Inter-lineage communication networks in keloid (CASE; right panel) and relatively normal skin tissues (CTRL; left panel). The total number of communications is shown for each cell lineage. The line color indicates that the ligands are broadcast by the cell lineage in the same color. The line thickness is proportional to the number of broadcast ligands. **(B)** Heatmap shows the number of communications between any two of lineages in CASE (right panel) and CTRL (left panel). **(C)** The ligand-receptor pairs shown significant changes in specificity between any one of the non-fibroblast lineages and fibroblasts in CASE versus CTRL. Fibroblasts express receptors and receive ligand signals from other lineages. The dot size reflects the P-value of the permutation tests for lineage-specificity. The dot color denotes the mean of the average ligand-receptor expression in the interacting lineages. **(D)** The ligand-receptor pairs shown significant changes in specificity between fibroblasts and any one of the non-fibroblast lineages in CASE versus CTRL. Fibroblasts express ligands and broadcast ligand signals for other lineages. FB: fibroblast; KTR: Keratinocyte; LEU: Leukocyte; lEndo: lymphatic endothelial cell; MLA: Melanocyte; SGC: sweat gland cell; SMC: smooth muscle cell; vEndo: vascular endothelial cell

### The heterogeneity and regulatory changes of vascular endothelial cells in keloid skin tissue

Keloid tissue is characterized by excessive capillary formation, and the dysregulation of vascular endothelial cells have been proposed to be associated with keloid progression (Tanaka et al., 2019). Therefore, we also examined the heterogeneity and regulatory changes of vascular endothelial cells in keloid skin tissue. Unbiased clustering revealed four clusters with distinct expression profiles (Figure 7A; Table S8), which could be marked with specific marker combinations: c2-*PECAM1*^+^ *HMOX1*^high^ CXCL3^-^, c4-*PECAM1*^+^ *ACKR1*^high^ *HMOX1*^-^, c5-*PECAM1*^+^ *CXCL12*^high^ *CXCL3*^-^ and c18-*PECAM1*^+^ *CXCL3*^+^ (Figure 7B). Correlation analysis revealed that c4 appears to be distant from the other clusters (Figure 7C). Functional enrichment analysis revealed that c4 represents antigen-presenting endothelial cells expressing the MHC class II markers such as *HLA-DRA* (Figure 7D), which has recently been reported (Han et al., 2020). Then, we identified the lineage-specific differentially regulated pathways of vascular endothelial cells in keloid versus normal skin tissue (Figure 7E; Figure S2; Table S9). As expected, vascular endothelial growth factor receptor (VEGFR) signaling pathway, which is important in pathological angiogenesis (Shibuya, 2011), was significantly activated. Intriguingly, tumor-related signaling pathways were activated for vascular endothelial cells in keloid, such as “oncogenic MAPK signaling”, “signaling by WNT in cancer” and “regulation of *PTEN* gene transcription”. In addition, we observed activated signaling pathways implicated in tumor angiogenesis, such as NOTCH signaling (NOTCH1/NOTCH4) (Dufraine et al., 2008), TGF beta receptor signaling (Pardali and Ten Dijke, 2009) and Eph-ephrin signaling (Cheng et al., 2002). Furthermore, we identified the ligand-receptor pairs altered in keloid between vascular endothelial cells and the other cell lineages (Figure 7F and 7G). Notably, EFNB2-EPHA4, a ligand-receptor pair in Eph-ephrin signaling, was significantly altered in keloid, which was well known for promoting sprouting angiogenesis (Kania and Klein, 2016). In keloid, the mean expression levels of EFNB2-EPHA4 between leukocytes, Schwann cells and vascular endothelial cells increased (Figure 7F), suggesting vascular endothelial cells were positively regulated via EFNB2-EPHA4 signaling. Meanwhile, vascular endothelial cells may also regulate the transcriptional states of fibroblasts and smooth muscle cells through EFNB2-EPHA4 signaling in keloid (Figure 7G). In addition, the VEGFB-FLT1(VEGFR1) interaction between leukocytes and vascular endothelial cells specifically occurred in keloid, reflecting the contribution of leukocytes to the activation of VEGFR signaling pathway in vascular endothelial cells (Figure 7F).

**Figure 7.**
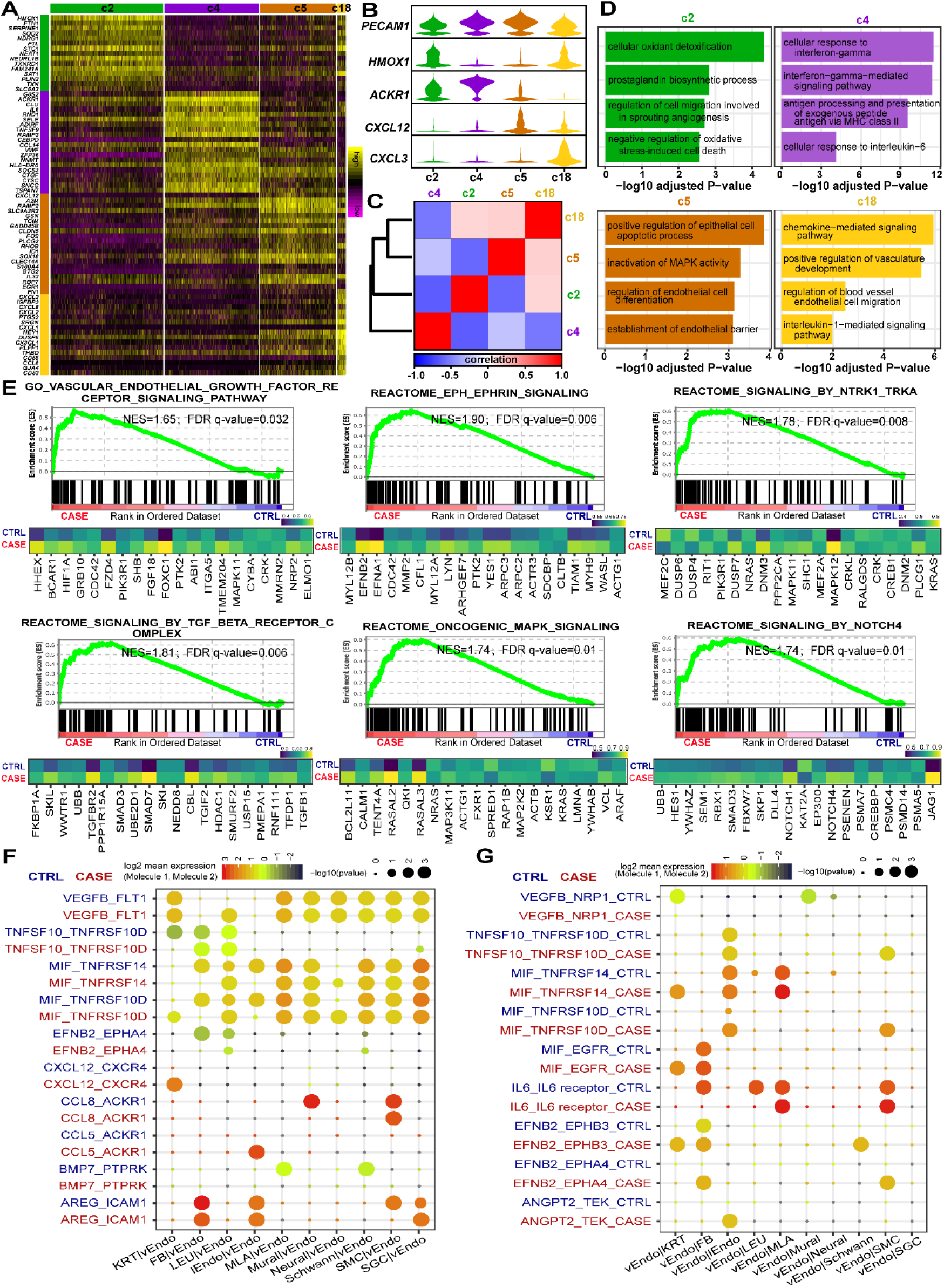
The heterogeneity and regulatory changes of vascular endothelial cells in keloid skin tissues. **(A)** Heatmap showing distinct expression profiles among the four vascular endothelial cell clusters. **(B)** Representative molecular signatures of the four cell clusters. **(C)** Correlations among the four cell clusters. **(D)** Functional enrichment for each of the four clusters. **(E)** Enrichment plots (upper panel) and leading-edge gene expression heatmaps (the top 20 genes; lower panel) for representative signaling pathways up-regulated in vascular endothelial cells in keloid tissues. NES: Normalized enrichment score, used to compare analysis results across gene sets. The vertical lines in the enrichment plot show where the members of the gene set appear in the ranked list of genes. Leading-edge genes: a subset of genes in the gene set that contribute most to the enrichment. Average expressions across cells in each group are shown in the heatmap. **(F)** The ligand-receptor pairs shown significant changes in specificity between any one of the non-vascular endothelial lineages and vascular endothelial cells in CASE versus CTRL. vascular endothelial cells express receptors and receive ligand signals from other lineages. The dot size reflects the P-value of the permutation tests for lineage-specificity. The dot color denotes the mean of the average ligand-receptor expression in the interacting lineages. **(G)** The ligand-receptor pairs shown significant changes in specificity between vascular endothelial cells and any one of the non-vascular endothelial lineages in CASE versus CTRL. Vascular endothelial express ligands and broadcast ligand signals for other lineages. FB: fibroblast; KTR: Keratinocyte; LEU: Leukocyte; lEndo: lymphatic endothelial cell; MLA: Melanocyte; SGC: sweat gland cell; SMC: smooth muscle cell; vEndo: vascular endothelial cell

## Discussion

Understanding the cellular heterogeneity and regulatory changes of tissues in diseased conditions is fundamental to successful medical therapy development. In this study, we performed single-cell RNA-seq of 28,064 cells from keloid skin tissue and adjacent relatively normal tissue. Unbiased clustering revealed substantial cellular heterogeneity of the keloid tissue, which included 21 cell clusters assigned to 11 cell lineages. Differential proportion analysis revealed significant expansion for fibroblasts and vascular endothelial cells in keloid compared with control, reflecting their strong association with keloid pathogenesis. Gene set enrichment analysis revealed fibroblast-specific dysregulated pathways in keloid versus normal tissue. Among them, signaling pathways implicated in keloid or other fibrotic disorders were found to be activated in keloid fibroblasts such as TGF beta, canonical WNT, PDGF, NOTCH1 and Eph-ephrin signaling pathways. Intriguingly, a set of genes involved in negative regulation of *PTEN* gene, a known tumor suppressor, were up-regulated in keloid fibroblasts. Previously unrecognized cellular heterogeneity of keloid fibroblasts was observed, which consists of five distinct subpopulations. Pseudo-temporal ordering of the fibroblasts reveals a branched trajectory with a significant shift towards myofibroblast phenotype in keloid. Branch-dependent genes that were putatively associated with fibroblast-to-myofibroblast phenotypic transition were identified. Comparative analysis of the gene regulatory networks of fibroblasts between keloid and normal tissue revealed dysregulated transcription factors and signaling receptors in keloid fibroblasts, which could serve as targets for medical interventions. Among them, *TWIST1, SMAD3* and *FOXO3*, regulators implicated in fibrogenesis, were both branch-dependent and dysregulated in the network of keloid fibroblasts. Notably, *EPHB2*, a receptor in Eph-ephrin signaling ranked at the top of the keloid network. Cell-cell communication analysis revealed a dense communication network among fibroblasts, vascular endothelial cells, neural cells and keratinocytes. Furthermore, ligand-receptor interaction changes specific to fibroblasts in keloid were identified. For example, the TGFB1-TGF beta receptor 2 interactions between non-fibroblast lineages and fibroblasts became significantly more specific in keloid. At last, we also examined the heterogeneity and regulatory changes of vascular endothelial cells in keloid tissue, which could be another targeted cell lineage for medical interventions besides of the fibroblasts. Four distinct subpopulations were identified. Intriguingly, tumor-related signaling pathways were activated for vascular endothelial cells in keloid, such as “oncogenic MAPK signaling”, “signaling by WNT in cancer” and “regulation of PTEN gene transcription”. Moreover, we identified the ligand-receptor pairs altered in keloid between vascular endothelial cells and the other cell lineages. Notably, EFNB2-EPHA4 and VEGFB-FLT1 interactions were significantly altered in keloid, which were known for promoting angiogenesis.

Single-cell analysis provided us a wealth of candidate molecules for developing targeted approaches to combatting the excessive ECM deposition caused by fibroblasts with aberrant proliferation and differentiation, or to suppressing active angiogenesis mediated by dysregulated vascular endothelial cells in keloid. Substantial evidence supports that TGF beta signaling pathway is central in keloid or other fibrotic disorders (Piersma et al., 2015). The ligand TGFB1 could reprogram fibroblasts to myofibroblasts, which synthesize large volumes of collagen fibers, a hallmark of keloid (Ashcroft et al., 2013; Nangole and Agak, 2019). *TWIST1* functions as a pro-fibrotic factor in a TGF beta/SMAD3/p38-dependent manner (Ning et al., 2018). In addition, TGF beta signaling has been implicated in tumor angiogenesis (Pardali and Ten Dijke, 2009). In this study, we found the activation of TGF beta receptor signaling in both fibroblasts (Figure 3B) and vascular endothelial cells (Figure 7E) from keloid tissue. *SMAD3*, the key downstream regulator of TGF beta signaling, was found to be critical for the dysregulated states of keloid fibroblasts through independent analyses (Figure 5D; Figure 5F). TGFB1-TGF beta receptor 2 interactions between non-fibroblast lineages and fibroblasts was found to become significantly more specific in keloid (Figure 6C). Together, these results consolidated our view that TGF beta signaling could serve as a medical target for simultaneously suppressing myofibrogenesis and angiogenesis in keloid. Similarly, Eph-ephrin signaling represents another promising target pathway to regulate the states of keloid fibroblasts and vascular endothelial cells. We found the activation of Eph-ephrin signaling in both fibroblasts (Figure 3B) and vascular endothelial cells (Figure 7E) from keloid tissue. We found *EPHB2*, a receptor in Eph-ephrin signaling, ranked at the top in the network of keloid fibroblasts (Figure 5E). The angiogenesis-promoting EFNB2-EPHA4 interactions (Kania and Klein, 2016) between vascular endothelial cells and others were significantly altered in keloid (Figure 7F and 7G). Besides of these two pathways, NOTCH, PDGF and VEGF signaling were also found to be probably important in keloid pathogenesis. Past research often focused on the isolated signaling cascades, whereas recent developments showed that parts of the signaling pathways may be organized into an intricate network (Piersma et al., 2015). So, it is possible that targeting anyone of these dysregulated pathways in keloid would be ultimately effective for preventing keloid formation or recurrence, although manageable side effects should be well considered in clinical trials.

Although keloid is generally regarded as a benign dermal tumor, keloid cells display some malignant features such as resistance to apoptosis, atypical differentiation and excessive proliferation (Lim et al., 2006). The mechanism underlying these features have not been fully elucidated. The *PTEN* gene encodes a negative regulator of PI3K/ACT/mTOR pathway that controls cell proliferation and survival (Chalhoub and Baker, 2009). The dysfunction of PTEN have been observed frequently in malignant tumors (a known tumor suppressor) and some fibrotic disorders such as lung fibrosis (Tian et al., 2019). Genetic studies revealed that polymorphisms in *PTEN* were associated with the increase keloid risk in Chinese Han population (Li et al., 2014). Reduced expression of *PTEN* has been observed in keloid tissue (Sang et al., 2015). We found the pathway of negatively regulation of *PTEN* transcription was activated in both keloid fibroblasts (Figure 3B) and vascular endothelial cells (Figure 7E). The genes implicated in epigenetic repression of *PTEN* were up-regulated in keloid, involving either the recruitment of the nucleosome remodeling and deacetylation (NuRD) protein complex (e.g., *HDAC1, HDAC3, HDAC7, CHD3* and *CHD4*), or the recruitment of the polycomb repressor complex (e.g., *CBX4, CBX8, EZH2, BMI1, MBD3* and *MECOM*). Together, these results reflect that the dysregulation of *PTEN*-mediated pathway may be responsible for the malignant features of keloid. In addition, other tumor-related signaling pathways such as “oncogenic MAPK signaling”, and “signaling by WNT in cancer” were activated for vascular endothelial cells in keloid (Figure 7E), which supports the view that overlap exists in the dysregulated pathways between keloid and malignant tumors. Our findings have implications for clinical treatment of keloid: medical therapies in tumor: for example, drugs targeting MAPK signaling in cancer (Lee et al., 2020), may also be effective in keloid treatment.

In conclusion, we provided the first systematic analysis of cellular heterogeneity and regulatory changes of keloid skin tissue at single-cell resolution. Our study put novel insights into the pathogenesis of keloid, and revealed dysregulated pathways in keloid fibroblasts and vascular endothelial cells, which could serve as potential targets for medical therapies. Our dataset constitutes a valuable resource for further investigations of the mechanism of keloid pathogenesis.

## Supplemental Materials

**Figure S1.** Differential proportional analysis of each cell lineages in CASE versus CTRL. A significant threshold of p-value < 0.05 for one-way paired t-tests was used.

**Figure S2.** Gene set enrichment analysis reveals lineage-specific dysregulated pathways for vascular endothelial cells in keloid versus normal skin tissue.

**Table S1.** Clinical information of the patients and sequencing quality metrics of the samples.

**Table S2.** Molecular signature for each of the 21 cellular clusters.

**Table S3.** Results of gene set enrichment analysis for the fibroblast lineage between CASE and CTRL (Sheet1: REACTOME; Sheet2: GENE ONTOLOGY).

**Table S4**. Molecular signature for each of the five subpopulations of the fibroblast lineage.

**Table S5.** Genes which expression changed as a function of the pseudotime inferred by Monocle2.

**Table S6.** Results of the node centrality comparisons between the gene regulatory networks of the fibroblasts in CASE and CTRL.

**Table S7.** Statistical inference of receptor-ligand specificity between all cell lineages with CellPhoneDB2.

**Table S8.** Molecular signature for each of the four subpopulations of the vascular endothelial cells.

**Table S9.** Results of gene set enrichment analysis for the vascular endothelial cells between CASE and CTRL (Sheet1: REACTOME; Sheet2: GENE ONTOLOGY).

## Methods

### Ethical approval

All human patient recruitments and tissue sampling procedure complied with the ethical regulations approved by Peking Union Medical College Hospital. Each subject received written informed consent.

### Sample preparation and tissue dissociation

Keloid lesional tissues were harvested during plastic surgery from four patients confirmed to have clinical evidence of keloid. No patient had received chemotherapy or radiotherapy prior to surgery. As a control, matched relatively normal skin tissues adjacent to the keloid scar were also sampled. Excised skin was immersed in the pre-cold Dulbecco’s Modified Eagle Medium (DMEM, CAT: 11965-084, Gibco™) culture medium with 10% FBS (CAT: 10270-106, Gibco™) and then transferred to the lab immediately. Enzymatic digestion and mechanical cutting were performed to dissociate the skin tissue to single-cell suspension. In brief, the skin tissue was washed twice with DMEM. After removal of the adipose tissue under the reticular dermis, the samples were minced into small pieces. Multiplex enzyme was used to digest the samples overnight according to the manufactory’s protocol. Next, mechanical cutting was performed on gentleMACS™ Octo Dissociator using ‘h_skin_01’ program. After that, fragments and large clumps were removed by filtering through a 100-μm filter (BD Falcon) and then a 40-μm filter. Dead Cell Removal Kit (Miltenyi Biotec) was used to remove the cells with low viability. The live cells were subsequently centrifuged at 300 relative centrifugal force (rcf) for 5 minutes at 4 °C to obtain a cell pellet, which was then diluted to 1×10E6 cells per millimeter.

### Single-cell RNA-seq library preparation and sequencing

Single-cell Gel Beads-in-Emulsion (GEM) generation, barcoding, post GEM-RT cleanup, cDNA amplification and cDNA library construction were performed using Chromium Single Cell 3’ Reagent Kit v3 chemistry (10X Genomics, USA) following the manufacturer’s protocol. The resulting libraries were sequenced on a NovaSeq 6000 System (Illumina, USA).

### Sample demultiplexing, barcode processing and UMI counting

The official software Cell Ranger v3.0.2 (https://support.10xgenomics.com) was applied for sample demultiplexing, barcode processing and unique molecular identifier (UMI) counting. Briefly, the raw base call files generated by the sequencers were demultiplexed into reads in FASTQ format using the ‘‘cellranger mkfastq’’ pipeline. Then, the reads were processed using the ‘‘cellranger count’’ pipeline to generate a gene-barcode matrix for each library. During this step, the reads were aligned to the mouse human reference genome (version: GRCh38). The resulting gene-cell UMI count matrices of all samples were ultimately concatenated into one matrix using the ‘‘cellranger aggr’’ pipeline.

### Data cleaning, normalization, feature selection, integration and scaling

The concatenated gene-cell barcode matrix was imported into Seurat v3.1.0 (Butler et al., 2018; Stuart et al., 2019) for data preprocessing. To exclude genes likely detected from random noise, we filtered out genes with counts in fewer than 3 cells. To exclude poor-quality cells that might have resulted from doublets or other technical noise, we filtered cell outliers (> third quartile + 1.5 × interquartile range or < first quartile - 1.5 × interquartile range) based on the number of expressed genes, the sum of UMI counts and the proportion of mitochondrial genes. To further remove doublets, we filtered out cells based on the predictions by Scrublet (Wolock et al., 2019). In addition, cells enriched in hemoglobin gene expression were considered red blood cells and were excluded from further analyses. The sum of the UMI counts for each cell was normalized to 10,000 and log-transformed. For each sample, 2,000 features (genes) were selected using the “FindVariableFeatures” function of Seurat under the default settings. To correct for potential batch effects and identify shared cell states across datasets, we integrated all the datasets via canonical correlation analysis (CCA) implemented in Seurat. To mitigate the effects of uninteresting sources of variation (e.g., cell cycle), we regressed out the mitochondrial gene proportion, UMI count, S phase score and G2M phase score (calculated by the “CellCycleScoring” function) with linear models using the “ScaleData” function. Then, the data were centered for each gene by subtracting the average expression of that gene across all cells, and were scaled by dividing the centered expression by the standard deviation.

### Dimensional reduction and clustering

The expression of the selected genes was subjected to linear dimensional reduction through principal component analysis (PCA). Then, the first 30 principal components of the PCA were used to compute a neighborhood graph of the cells. The neighborhood graph was ultimately embedded in two-dimensional space using the non-linear dimensional reduction method of uniform manifold approximation and projection (UMAP) (Becht et al., 2019). The neighborhood graph of cells was clustered using Louvain clustering (resolution=0.6) (Blondel et al., 2008).

### Differential expression and function enrichment analysis

Differentially expressed genes between two groups of cells were detected with the likelihood-ratio test (test.use: ‘‘bimod’’) implemented in the ‘‘FindMarkers’’ function of Seurat. The significance threshold was set to an adjusted P-value < 0.05 and a log2-fold change > 0.25. Functional enrichment analyses of a list of genes were performed using ClueGO (Bindea et al., 2009) with an adjusted P-value threshold of 0.05.

### Gene set enrichment analysis

All the expressed genes were pre-ranked by Signal2Noise (the difference of means between CASE and CTRL scaled by the standard deviation). Then, the ranked gene list was imported into the software GSEA (version: 4.0.1) (Subramanian et al., 2005). An FDR q-value < 0.05 was considered to be statistically significant. Pre-compiled gene sets including REACTOME pathways and GENE ONTOLOGY biological processes in MSigDB (version: 7.0) (Liberzon et al., 2015) were used in this analysis. The results were visualized using the EnrichmentMap plugin of Cytoscape.

### Pseudo-temporal ordering of single cells

Pseudo-temporal ordering of the cells along the differentiation trajectory was performed using Monocle2 (Qiu et al., 2017). Briefly, the ordering was based on 1,000 genes that differed in expression between clusters selected via an unsupervised procedure: “dpFeature”. Then, the data space was reduced to two dimensions with the method “DDRTree”. The cells were ultimately ordered in pseudotime, and cells exhibiting high expression of myofibroblast markers were considered to represent the end of the trajectory.

To find significantly branch-dependent genes in their expression, we used the test named branched expression analysis modeling (BEAM), and the statistically significant threshold was set to a q-value < 1E-04.

### Gene regulatory network analysis based on single-cell transcriptomes

Gene regulatory networks were constructed from single-cell datasets and compared using the method implemented in bigScale2 (Iacono et al., 2019). Briefly, gene regulatory networks for CASE and CTRL were inferred with the ‘compute.network’ function (clustering=‘direct’, quantile.p = 0.90) separately. Genes encoding ribosomal proteins or mitochondrial proteins were excluded from this analysis. Then, the number of edges were homogenized throughout the obtained networks using the ‘homogenize.networks’ function. Finally, changes in node centralities (the relative importance of genes in the network) in CASE compared to CTRL were identified using the the ‘compare.centrality’ function. Four measures of centrality, namely degree, betweenness, closeness and pagerank, were considered. The networks were ultimately visualized with Cytoscape (version: 3.7.0).

### Cell-cell communication analysis based on single-cell transcriptomes

To analyze cell-cell communication based on single-cell transcriptomic datasets, we used CellPhoneDB 2.0 (Efremova et al., 2019), which contains a curated repository of ligand-receptor interactions and a statistical framework for inferring lineage-specific interactions. Briefly, potential ligand-receptor interactions were established based on expression of a receptor by one lineage and a ligand by another. Only ligands and receptors expressed in greater than 10% of the cells in any given lineage were considered. The labels of all cells were randomly permuted 1000 times and the means of the average ligand-receptor expression in the interacting lineages were calculated, thus generating a null distribution for each ligand-receptor pair in each pairwise comparison between lineages. Ultimately, a p-value for the likelihood of lineage specificity for a given ligand-receptor pair was obtained.

## Author contributions

Liu analyzed the data, interpreted the results and wrote the manuscript. W. C. performed tissue dissociation and library preparation, and participated in drafting the manuscript. M., Z. L., T. M., J. C. and N. Y. prepared the samples and contributed to the result interpretation. X. Long and Z. Z. conceived the project. X. Liu and W. C. also participated in the design of the project.

## Acknowledgments

This work was supported by the grants from the Natural Science Foundation of China (81870229,81900282) and the Chinese Academy of Medical Sciences Initiative for Innovative Medicine Grant (2016-I2M-1-016).

