## Supplementary figures and images for "Single-cell RNA-seq reveals lineage-specific regulatory changes of fibroblasts and vascular endothelial cells in keloid"

### Figure S1

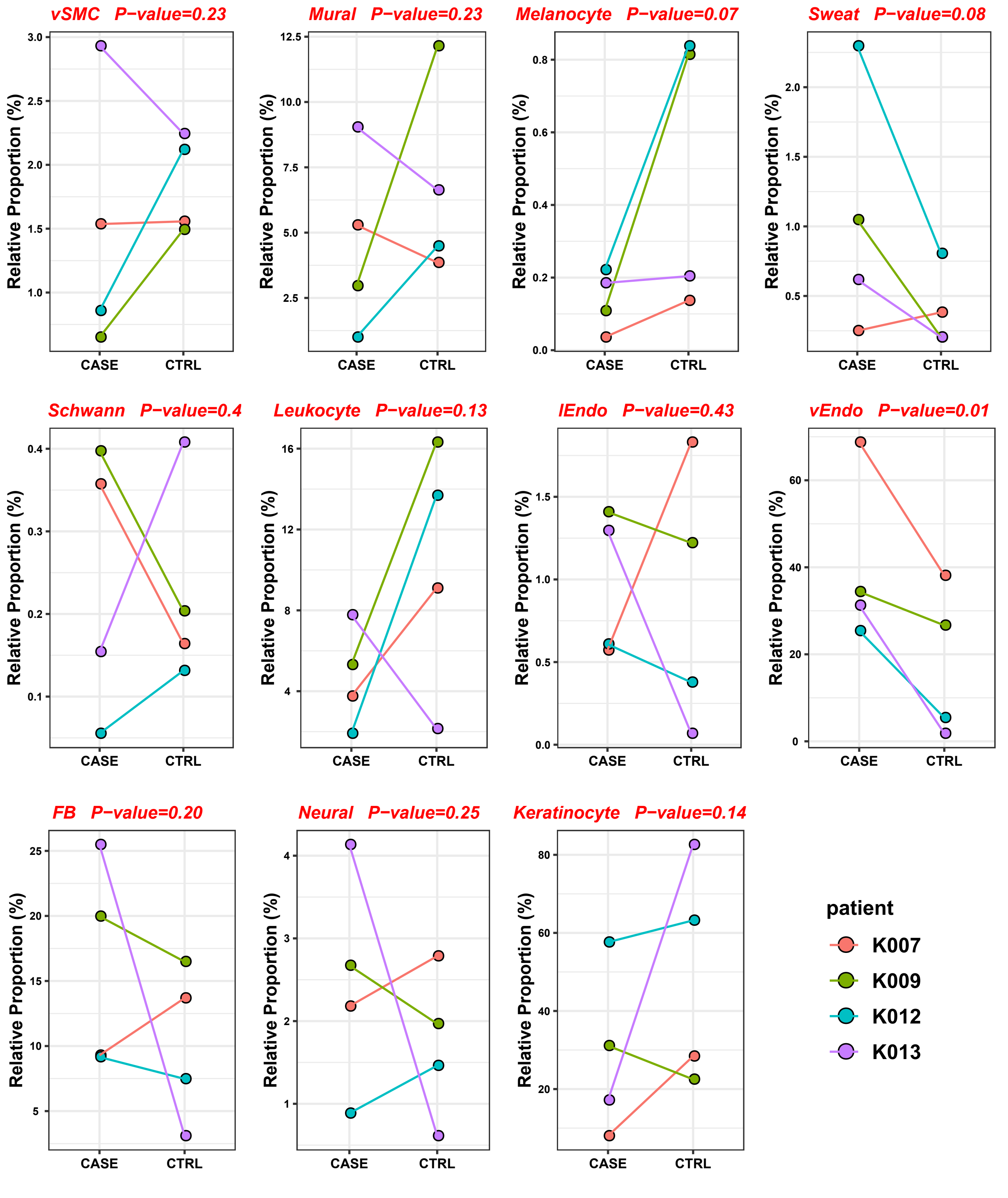

### Figure S2

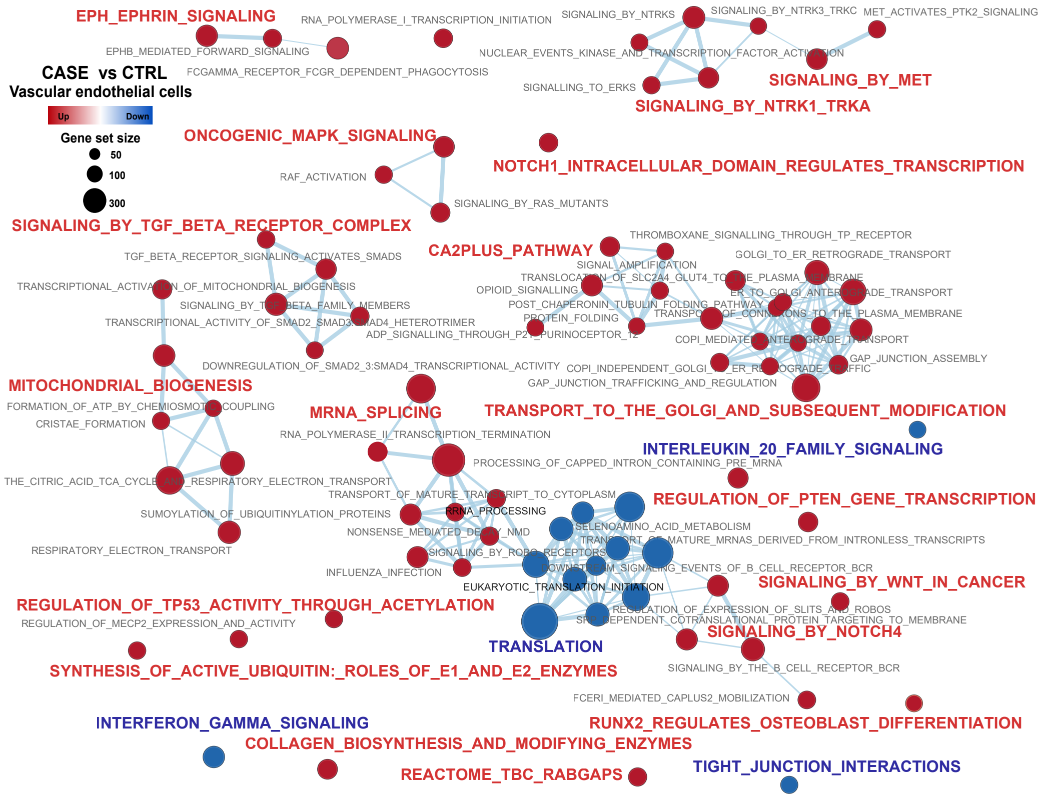
